# Rubisco forms a lattice inside alpha-carboxysomes

**DOI:** 10.1101/2022.01.24.477598

**Authors:** Lauren Ann Metskas, Davi Ortega, Luke M. Oltrogge, Cecilia Blikstad, Tom Laughlin, David F. Savage, Grant J. Jensen

## Abstract

Bacteria employ microcompartments to sequester enzymatic processes, either for purposes of protecting cellular contents from reactive intermediates or as a way of increasing reaction efficiency. In these structures, a cargo of enzymes and accessory proteins is encased within a semi-permeable protein shell that permits passage of substrates and products but restricts movement of intermediates. In addition to their importance as a component of many bacterial species’ metabolisms, microcompartments have recently become a target of protein engineering. The shells can be reassembled from purified proteins, and the full operons can be functionally expressed outside their native prokaryotes and can remain functional following purification. Despite the importance of microcompartments in prokaryotic biology and bioengineering, structural heterogeneity has prevented a complete understanding of their architecture, ultrastructure, and spatial organization. Here, we employ cryo electron tomography to image α-carboxysomes, a pseudo-icosahedral microcompartment responsible for carbon fixation. We have solved a high-resolution subtomogram average of the Rubisco cargo in situ, and determined a novel arrangement of the enzyme. We find that the H. neapolitanus Rubisco polymerizes in vivo, mediated by the small Rubisco subunit. These fibrils can further pack to form a lattice with six-fold pseudo-symmetry. This arrangement preserves freedom of motion and accessibility around the Rubisco active site and the binding sites for two other carboxysome proteins, CsoSCA (a carbonic anhydrase) and the disordered CsoS2, even at Rubisco concentrations exceeding 800 μM. This characterization of Rubisco cargo inside the α-carboxysome provides new insight into the balance between order and disorder in microcompartment organization.

## INTRODUCTION

The enzyme Rubisco is the primary catalyst for biological carbon dioxide fixation.^1^ In many autotrophic bacteria, Rubisco is sequestered inside carboxysomes, proteinaceous microcompartments that concentrate carbon dioxide to improve turnover of the enzyme.^2,3^ While many of the individual proteins are structurally characterized, the supramolecular assembly of the entire carboxysome complex remains less well understood. Here, we use cryo electron tomography and subtomogram averaging to determine an *in situ* structure of Rubisco at 4.5 Å, as well as identify the position and orientation of each Rubisco complex within the carboxysome. Using these data, we find that Rubisco polymerizes in the highly concentrated carboxysome interior and that these Rubisco fibrils can further pack into a twisted hexagonal lattice. These findings give insight into how bacteria have evolved their use of the most abundant enzyme on earth to optimize the efficiency of carbon fixation and provide future directions for bioengineering of microcompartments.

Many prokaryotes employ compartmentalization to sequester and facilitate biochemical activities. One example of this organization is the bacterial microcompartment, a collection of enzymes enclosed in a proteinaceous shell that selectively restricts passage of key intermediates and improves on-target catalysis.^4,5^ Interest in microcompartment bioengineering has recently grown, particularly for transplanting or reconstructing carbon fixing microcompartments to non-native hosts.^6–9^ However, while reconstituted assemblies of shell proteins and other components have been studied at high resolution,^6,10–12^ the supramolecular organization of the microcompartment interior remains elusive.

The α-carboxysome (CB) of *Halothiobacillus neapolitanus* is a microcompartment responsible for carbon fixation in cyanobacteria and some chemoautotrophs.^13^ It contains two enzymes, Rubisco and carbonic anhydrase, encapsulated within a pseudo-icosahedral protein shell (Figure 1).^14^ A third abundant component, CsoS2, is a disordered protein known to bind both Rubisco and shell *in vitro*, ^15,16^ and could potentially be involved in the clustering of Rubisco observed in the cytoplasm during CB assembly.^16–18^

**Figure 1.**
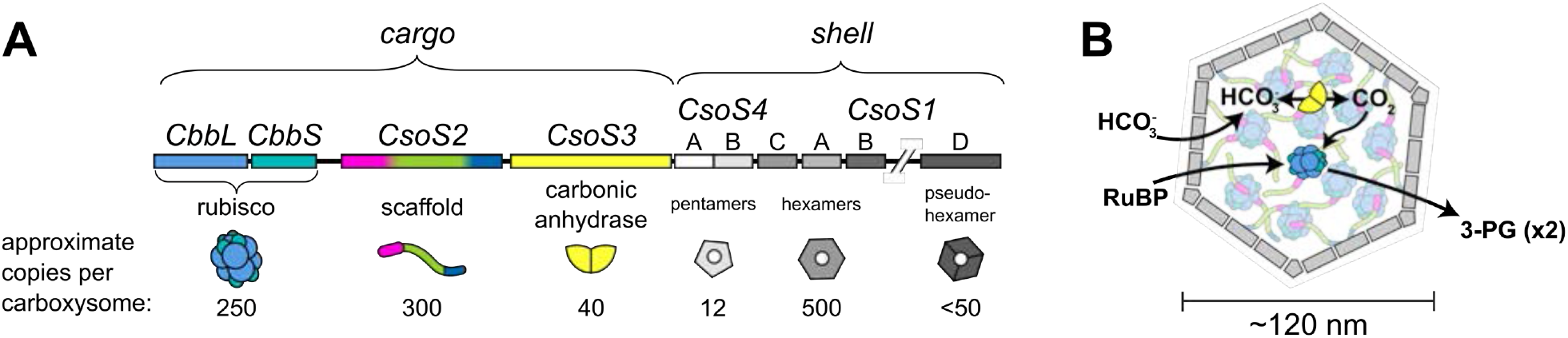
A: the *H. neapolitanus* CB operon, with estimates of protein copy numbers per carboxysome.^13^ B: current working model of a carboxysome. Carbonic anhydrase converts bicarbonate to carbon dioxide; Rubisco then uses the carbon dioxide to convert one molecule of ribulose-1,5-biphosphate to two 3-phosphoglycerate molecules.

Rubisco is a famously inefficient enzyme, with slow turnover and an undesirable off-pathway reaction with oxygen.^19^ However, encapsulation of Rubisco within the CB increases the organism’s total CO_2_ fixation rate.^2^ This is believed to be the result of a high local CO_2_ substrate concentration, which results from a non-equilibrium bicarbonate pool that is converted to CO_2_ inside the CB and restricted from diffusing away by a semi-permeable CB shell.^2,3^ Localized CO2 maximizes turnover and outcompetes oxygenase activity. However, it is unknown whether encapsulation of Rubisco in a CO_2_-rich environment is sufficient to explain the increased turnover, or if an additional feature such as enzymatic scaffolding or allosterically-regulated kinetics may also be required.

Here, we present a high-resolution view of Rubisco structure and organization within the *H. neapolitanus* carboxysome. We discover that Rubisco can arrange into a low-periodicity lattice inside the carboxysome at high concentration, driven by polymerization of the Rubisco itself. This unique feature of the α-carboxysome likely helps to increase the rate of carbon fixation inside the CB.

## RESULTS

We collected cryo electron tomograms of 139 purified α-carboxysomes displaying a range of sizes and Rubisco concentrations (Figure 2A, Extended Data Figure 1). Using a customized particle-picking approach designed to pick every Rubisco within the CBs (Extended Methods), we identified 32,930 Rubisco particles and carried out subtomogram alignment and averaging.

**Figure 2:**
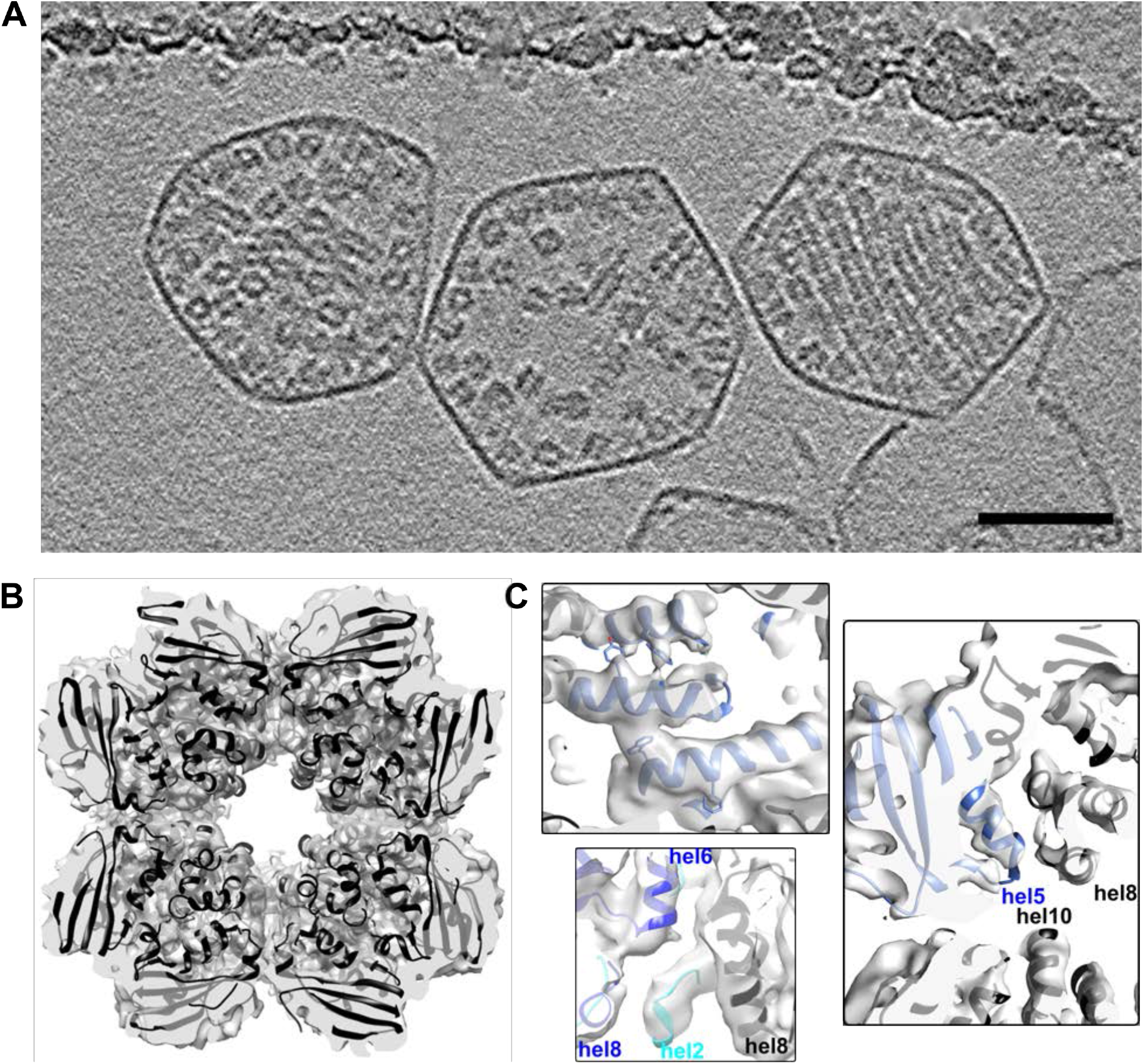
Rubisco subtomogram averaging. A) 4.4 nm-thick orthoslice through three carboxysomes with different packing behaviors. From left: dense, sparse, ordered. Scale bar 50 nm. B) Slab view of the 4.5 Å density map, with the 1SVD crystal structure docked inside (ribbon). C) 1SVD crystal structure docked within the density map (blue, large subunit; cyan, small subunit; black, neighboring subunits). Top: some bulky side chains (Y244, W276, W207, F213) are visible in the map. Right: helices are resolved and the docked crystal structure fits well across three large subunits. Bottom: the small subunit docks well between two large subunits.

Our 4.5 Å *in situ* Rubisco map shows strong agreement with crystal structure 1SVD of the same enzyme (Figure 2B). Helices align well to predicted locations in the crystal structure, and bulky side chain densities often match crystal structure positions (Figure 2C). However, we observe a modest discrepancy between our density map and the crystal structure in the small Rubisco subunit. If the large and small Rubisco subunits are docked as rigid bodies, this results in a slight (1 Å displacement) tilt of the beta-turn-beta feature of the small subunit (Y96 and adjacent residues, Extended Data Figure 2). This slightly widens the recently identified binding pocket for carboxysomal proteins CsoS2 and CsoSCA,^15,20^ but leaves the top and bottom interfaces of the complex unchanged relative to the crystal structure.

Roughly one third of the carboxysomes displayed ordered packing, with high concentrations of Rubisco packed tightly into aligned rows (Figure 2A). Other CBs displayed a range of Rubisco concentrations, from sparse to dense (Extended Data Figure 1). A nearest-neighbor Rubisco alignment search across all CBs revealed a concentration dependence: Rubisco orientation is random at low concentration (sparse CBs), but becomes increasingly non-random as concentration rises (dense CBs, Extended Data Figure 3). To differentiate between dense and ordered CBs, we used the second rank order tensor to quantify the alignment of all the Rubiscos within a CB (Figure 3A, Extended Methods).^21^ Dense CBs have localized Rubisco alignment, while ordered CBs show global Rubisco alignment (S > 0.35). Ordered CBs are only found with Rubisco concentrations above 650 μM (Figure 3A).

**Figure 3:**
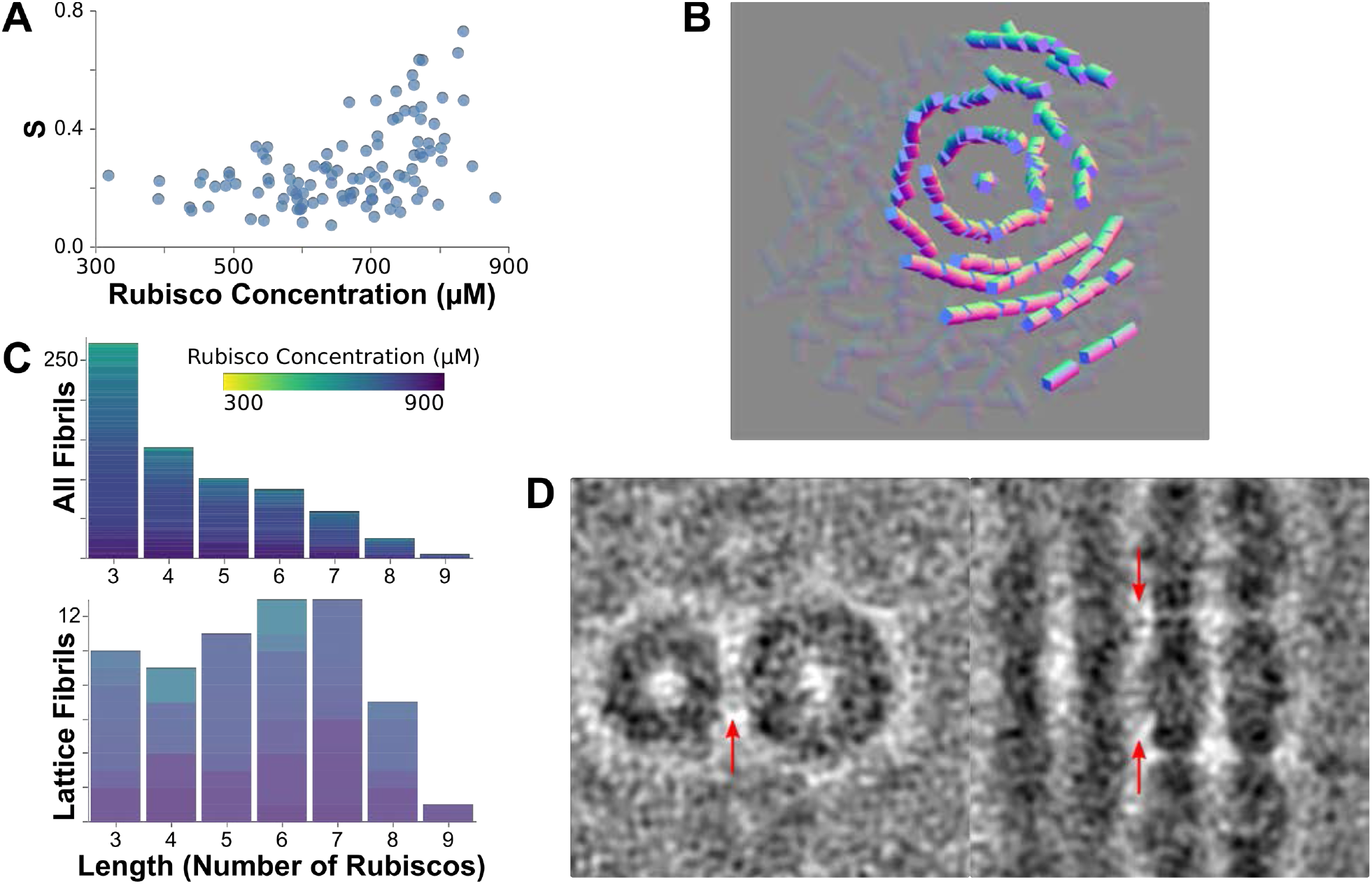
Rubisco forms a twisted hexagonal lattice. A) The second rank-order scalar S indicates global alignment along one axis in ordered CBs at concentrations exceeding 650 μM. B) A representative rendering of one Rubisco lattice shows a central fibril of Rubisco with six angled fibrils packed around it. C) Histograms of fibril length for all fibrils (upper) and inner-lattice fibrils (lower) show that lattice formation is associated with increased fibril length. D) Subtomogram averages show loose registration between the fibrils. Faint density in the fibril interface (red arrows) does not average to a specific interaction site.

All ordered CBs contain both nematic and isotropic phases in different areas of the interior. The short fibril lengths and disorder in the CB preclude traditional analyses for lattices and liquid crystals, but fibril subunits can be identified through spatial analysis (Extended Methods). Rendering all Rubisco fibrils within a CB displays the ordered phase: a loose hexagonal lattice-like ultrastructure of Rubisco fibrils twisting about each other with six-fold pseudo-symmetry (Figure 3B). The fibrils are held at a distance of 12.5 ± 0.7 nm with a tilt of 10 ± 3 degrees (Figure 3D).

Despite this order, no rigid scaffold is observed holding the Rubisco lattice together. An average of adjacent fibrils shows faint densities between the large and small subunits of adjacent Rubiscos (Figure 3D). We hypothesize that a disordered, lower-occupancy binding partner may maintain a maximum distance and promote tilt between fibrils. Such a linker would likely not be visible in a subtomogram average due to the combination of low occupancy and disorder, especially if there were also disorder in the binding site, causing the system to act as a ‘fuzzy complex’.^22^ CsoS2 has the appropriate disorder, length and Rubisco binding sites to serve this role.^15^

The fibrils forming the lattice appear to be the product of Rubisco polymerization. The ratio of bound and free Rubisco concentrations scales with Rubisco concentration in the CB (Figure 4A), and a difference map of Rubisco monomer removed from a fibril average shows no additional density except for the other Rubiscos (Figure 4C). Polymerization is observed in all packing types, though fibril lengths are longer inside the Rubisco lattices of ordered CBs (Figure 3C). This is likely the effect of oriented macromolecular crowding restricting diffusion away from the binding site and encouraging rebinding following dissociation.

**Figure 4:**
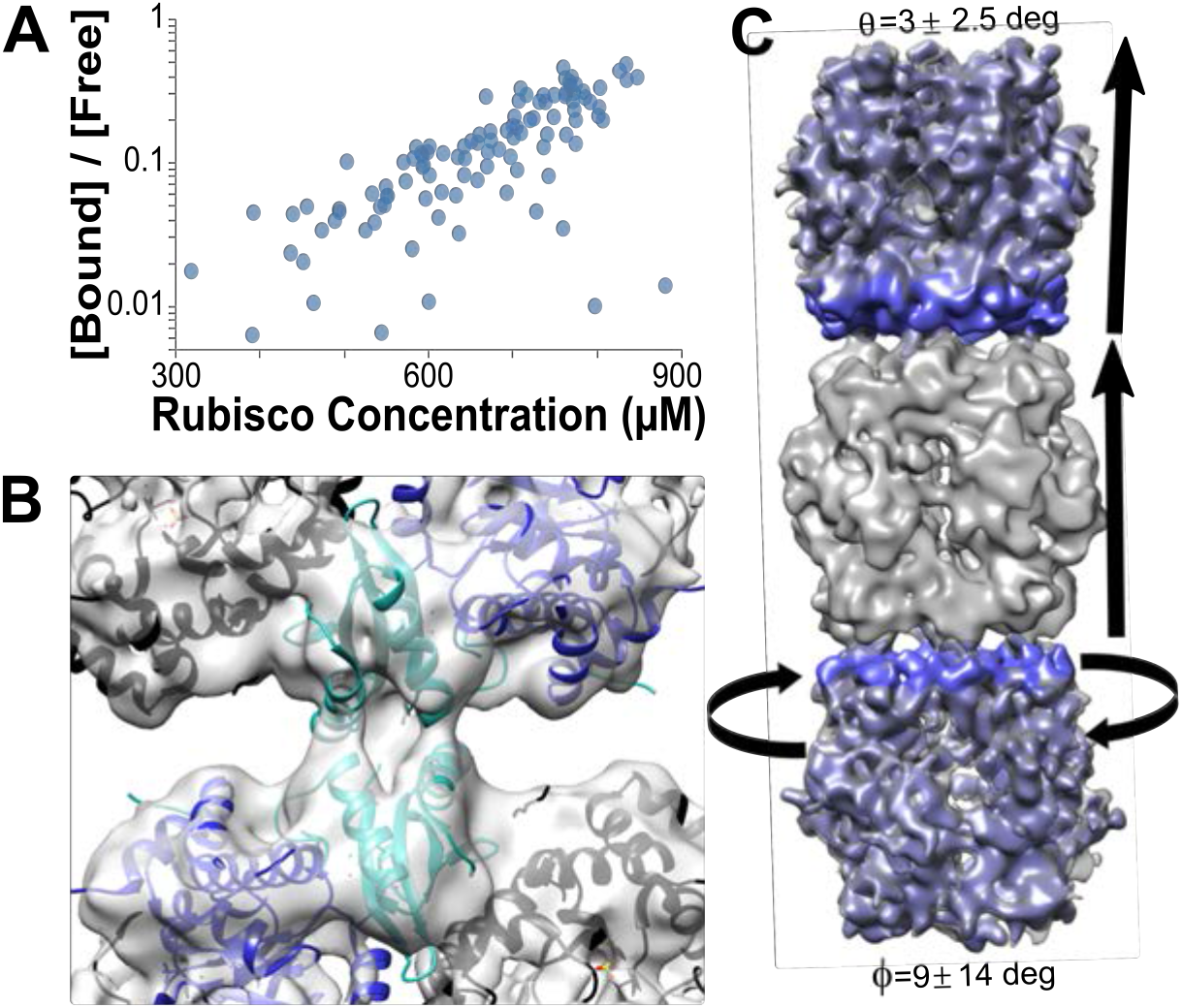
Rubisco polymerizes to form fibrils. A) Rubisco-Rubisco binding scales with concentration. B) A 10 Å slab through the subtomogram average of the Rubisco-Rubisco interface with the docked 1SVD crystal structure shows an interaction between helices 1 and 3 of the small subunit (cyan; large subunit, blue; additional subunits, black). C) A subtomogram average of Rubisco fibrils (grey) shows variable tilt and twist in the interaction. A difference map of the fibril and free Rubiscos shows density only in the Rubiscos above and below (blue), with no other protein mediating the interaction.

The Rubisco-Rubisco interaction appears to be low affinity, as the four binding sites on the C4 planar surface are not fully bound at any time. There is a bending angle of 3 ± 2.5 degrees and a 14-degree standard deviation in the twist between stacked Rubiscos, and non-symmetrized subtomogram averages of the fibrils have incomplete occupancy across the sites. It is likely that the low affinity of the Rubisco-Rubisco interaction results from a fast *k*_off_ rate at each site, decreasing the likelihood of all sites being bound at any one time and resulting in the wobbly interaction observed. This is consistent with the high concentrations associated with fibril formation.

We identified the Rubisco-Rubisco interaction site by performing subtomogram alignment and averaging of Rubisco within fibrils, using a tight mask at the upper interface to counteract the non-zero bending angle. The strongest density is between the tip of small subunit helix 1 on one Rubisco and the center of small subunit helix 3 of the binding partner, though the resolution is too poor to resolve the side chains driving the interaction. The variability in fibril parameters suggests that multiple loose interactions are likely responsible for the binding, consistent with the poorer resolution in the Rubisco-Rubisco interface.

Rubisco crystal structures also display variable interactions in this region. *H. neapolitanus* Rubisco structures 1SVD and 6UEW both show alternative longitudinal interactions in the crystal structure involving helix 3 (Extended Data Figure 4). Both crystal structure interactions lie within the range represented in our data, suggesting these conformations may be sampled *in vivo* (though not the dominant conformation). The side chains observed in the interaction site of the crystal structures do not have obvious complementarity. Taken together, these observations suggest that crystal packing may promote a low-affinity interaction with high symmetry and periodicity, one which is possible at the Rubisco concentrations present in the CB.

The Rubiscos adjacent to the shell occasionally participate in the long fibrils, but not consistently, and appear to behave differently than Rubisco in the CB interior. The number of Rubiscos in this shell-proximal layer scales with carboxysome volume (Figure 5A), and the layer is well-populated even in otherwise sparse CBs. We analyzed the angle of the Rubisco C4 axis relative to the shell and found a predominantly random distribution with a skew toward perpendicular orientations (likely reflecting the participation in fibrils, which would favor this orientation) (Figure 5B).

**Figure 5:**
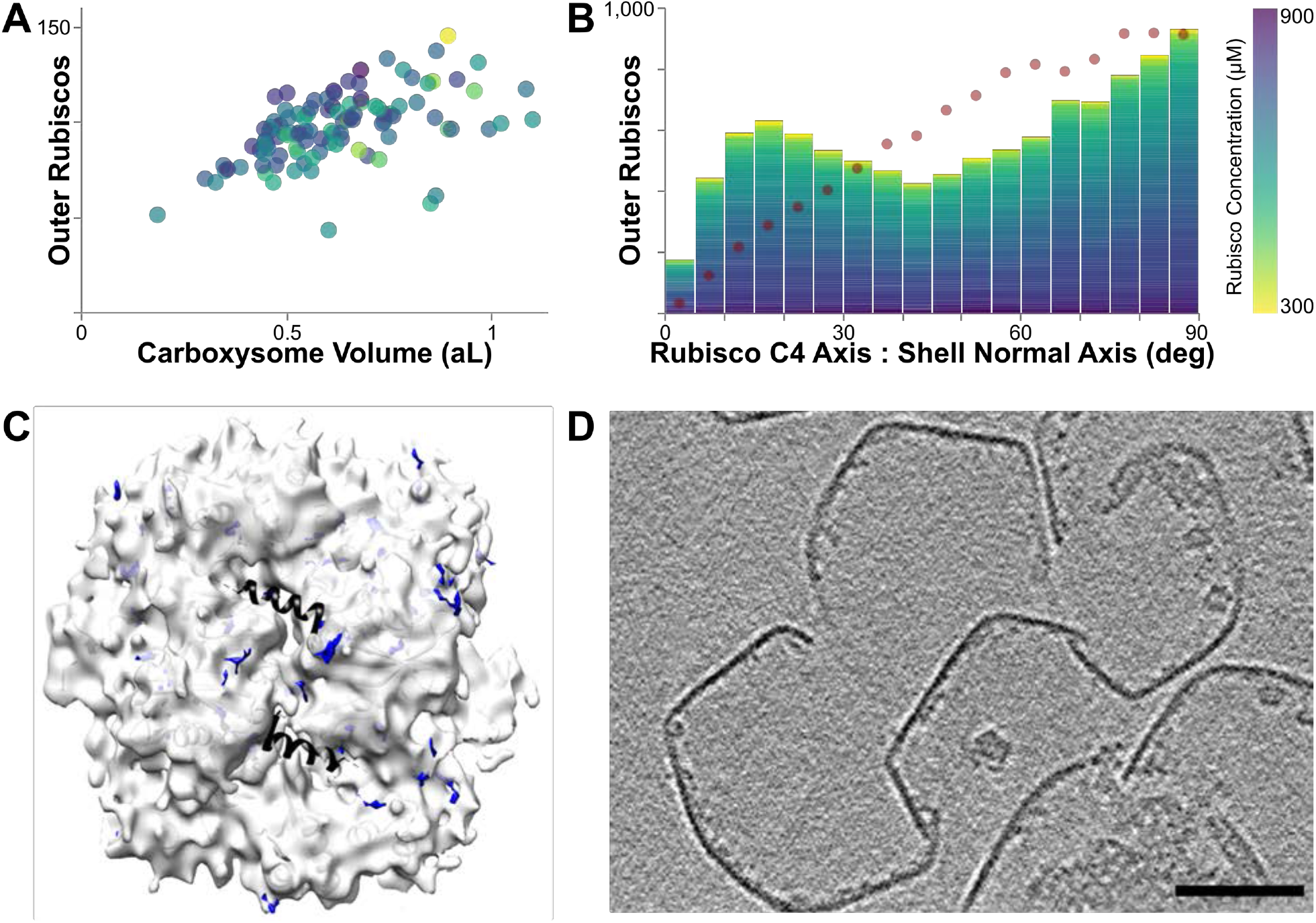
Rubisco in the outer layer. A) Rubisco lines the shell in a volume-dependent but concentration-independent manner. B) A histogram of the angle between the Rubisco C4 axis and the normal axis of the adjacent shell shows a modest preference for perpendicular orientations. Dotted line indicates random orientation. C) We do not observe strong CsoS2 occupancy in the outer Rubisco layer. White: 7 Å C1 interior Rubisco subtomogram average (3 nm slab). Blue: difference map showing density added in the outer layer Rubisco subtomogram average. Black: 6UEW crystal structure, CsoS2 peptide. D) A 4.4 nm-thick orthoslice of shell-associated density in broken shells. Scale bar 50 nm.

We were unable to identify any physical tether between Rubisco and the shell. We compared a subtomogram average of shell-adjacent Rubisco with an average of interior non-fibril Rubisco and found no density differences nor substantial density at the binding site for CsoS2 (Figure 5C). We did however observe many shell-attached densities in the tomograms, which are best visible in tomograms of broken CB shells (Figure 5D). Some of these densities are the correct size for carbonic anhydrase, which also is capable of binding Rubisco and was previously found to be shell associated.^20,23^ Protein identity could not be confirmed due to the enzyme’s small size.

The most consistent feature of the CBs is their compositional and structural heterogeneity. Subpopulations of CBs occasionally contain what appears to be an aggregate of polypeptide, presumably CsoS2; roughly 5% of CBs also contain a large enzyme that is consistent in size and shape with a 20S proteasome (Extended Data Figure 5). One CB had fibrils arranged parallel to the shell in concentric layers, rather than the hexagonal twisting ultrastructure observed in the other CBs (Extended Data Figure 5). Finally, even in the most ordered α-carboxysomes, a substantial portion of the Rubisco does not participate in the lattice (Figure 3B). This heterogeneity is consistent with models of CB assembly, which are based on phase separation and binding affinities rather than a tightly ordered, regular assembly mechanism.^15,17,24^

## DISCUSSION

Previous carboxysomal tomography studies used radial averaging to show that Rubisco formed a shell-adjacent layer,^14,17^ but the resolution in these studies was not high enough to determine Rubisco orientations necessary for ultrastructure analysis. Recently, tomographic analysis of the algal pyrenoid has shown that Rubisco behaves in a liquid-like fashion.^12,25,26^ In contrast, our high resolution analysis of organization in the α-carboxysome indicates that Rubisco organizes into a lattice-like ultrastructure in some CBs.

We observed three Rubisco packing types in the CBs: sparse, dense and ordered (Figure 2A). Sparse carboxysomes lack the concentration necessary to form an ultrastructure, but the dense and ordered packings overlap in Rubisco concentration. All packing types display polymerization of the Rubisco; therefore, the primary distinction between dense and ordered CB packing is whether the fibrils align, which supports fibril elongation and expansion of the nematic phase. In the β-CB system, characterization of reconstituted components and whole particles indicates that the interior becomes oxidized during CB maturation, with a concomitant increase in the mobility of interior cargo proteins.^12,24,27^ There is biochemical evidence that the α-CB may mature through a similar oxidation process.^28^ Therefore, the phase transition between dense and ordered packing may involve the CB maturation trajectory.

Rubisco fibril formation is concentration-dependent. Polymerization is a highly effective packing strategy, and does not obstruct the Rubisco active site nor CsoS2 and CsoSCA binding sites.^15^ However, symmetric protein complexes have an innate tendency toward long fibril formation,^29^ and uncontrolled cytoplasmic polymerization of the Rubisco could be deleterious and would likely disrupt CB formation.^17^ Therefore, the low affinity is likely tuned to ensure that polymerization occurs only when the local Rubisco concentration is increased by CsoS2 phase separation and then secured inside the CB shell. It is possible that the Rubisco polymerization may assist in CsoS2-based Rubisco condensation during carboxysome shell assembly.

The six-fold symmetry in the Rubisco fibril packing contrasts with the four-fold symmetry of the Rubisco itself. The large standard deviation in the fibril twist obscures the lateral face of the fibril, likely facilitating this symmetry break. This is in contrast to Form I Rubisco crystal structures in the PDB, which often have four-fold symmetric packing of fully parallel and untwisted fibrils. This four-fold Rubisco crystal packing is likely selected against *in vivo* because it could interfere with Rubisco function by obstructing motion of loop 6 during the catalytic cycle,^30^ obscuring peptide binding sites, and limiting diffusion of substrates and products. In contrast, the six-fold fibril arrangement has at least 1 nm of space around the circumference of each enzyme. The six-fold lattice may therefore be a packing mechanism to preserve function at concentrations that could otherwise crystallize or sterically impede function.

The higher-order arrangement of enzymes is often a component of their function and regulation.^31–33^ Rubisco itself has a rich representation in complex ultrastructures, from plant chloroplasts to algal pyrenoids to bacterial carboxysomes. The Rubisco ultrastructure within the *H. neapolitanus* carboxysome is novel in its design of loosely structured polymers rather than either a liquid-like encapsulation or a highly regular ultrastructure. Rubisco thus provides a unique comparative system for investigating the physical organization of the cell. Future studies will be needed to further elucidate the mechanisms underlying the observed divergent ultrastructural arrangements and their role in the control and regulation of this an important enzyme.

## AUTHOR CONTRIBUTIONS

CB and TL purified the carboxysomes. LAM performed cryo electron tomography and subtomogram averaging. LAM, DO, and LO analyzed the data. All authors designed research, interpreted results, and wrote the manuscript.

## ACKNOWLEDGEMENTS

Cryo electron microscopy was done in the Beckman Institute Resource Center for Transmission Electron Microscopy at Caltech, and subtomogram alignment and averaging used the Caltech Resnick High Performance Computing Center. We thank S. Chen and A. Malyutin for assistance with tomography data collection, and A. Burt for pseudocode to transition between Dynamo and Relion software packages. This work was supported by a Ruth L. Kirschstein NRSA Individual Postdoctoral Fellowship F32 1F32GM135994-01 to LAM, NIH R01GM129241 to DFS, and NIH R01 AI127401 to GJJ.

The authors declare no competing interests.

## METHODS

### Carboxysome Preparation

α-carboxysomes from *H. neapolitanus* were purified as described previously.^9^ Briefly, wild type *H. neapolitanus* cells were grown in DSMZ-68 medium in a 10 L chemostat. A ∼8 g cell pellet (resulting from 10-15 L of culture) was chemically lysed and purified by centrifugations and sucrose gradient. Quality and purity of sample was analyzed by Coomassie stained SDS-PAGE and negative stain TEM. The final sample had an A280 of roughly 10, and was stored at 4 °C in TEMB buffer (10 mM Tris pH 8, 1 mM EDTA, 10 mM MgCl_2_, 20 mM NaHCO_3_) until use.

### Tomography

10 nm gold colloids were mixed with 5% BSA in PBS to passivate the surface, then were buffer-exchanged into TEMB buffer and concentrated. The CB sample was diluted with the BSA-coated gold and plunge-frozen onto glow-discharged C-flat 2/2-300 EM grids using a Vitrobot. Imaging was performed on a Titan Krios with a Gatan K3 camera and energy filter. Tomograms were collected in SerialEM using a 1.104 Å pixel size (nominal magnification of 81,000) and a dose-symmetric tilt scheme moving from −66 to +66 degrees in 3-degree increments.^34,35^ Nominal defocus was varied across −1 to −5.5 μm for collection. Exposure was 2.9 electrons per Å^2^ per tilt, for a total dose of roughly 130 electrons. Frame alignment, 2D image processing, and tilt alignment were performed in etomo; CTF estimation in ctffind4; and tomogram reconstruction used novaCTF with the phaseflip option.^36–38^ Dose weighting was performed after defocus estimation but before tilt alignment.^39^

### Subtomogram Averaging

Particles were identified in SIRT-like filtered tomograms using a customized workflow designed to identify every Rubisco complex within each CB. The center of the CB shell was manually segmented in the xz and xy planes in IMOD,^40^ and a three-dimensional grid of Dynamo model points spaced 30 bin2 pixels apart was generated to fill this space using the Matlab inpolygon function. One iteration of subtomogram alignment was performed in Dynamo in bin4 against a reference average of Rubisco complexes outside the CB, with D4 symmetry imposed and translation searches limited.^41^ Duplicate positions were removed, then the revised model positions were sent to Relion for extraction and classification.^42^ Out of four classes, Relion consistently grouped subtomograms as Rubisco, shell or noise. This approach yielded Rubisco particles with an estimated 15-20% false positive and false negative rates. Classified particles were manually screened using a rule-based approach, eliminating Rubisco identifications outside the shell, shell identifications on Rubisco fibrils, identifications too close together, and so on. Screening was performed conservatively.

Following manual screening, the estimated false negative and false positive rates fell to 1-2% of total Rubisco particles within a CB when compared with manual identification in a clear, high-defocus tomogram. Unfiltered map resolutions were not degraded by using every single particle identified (Extended Data Figure 6), further strengthening the evidence for the strong success rates in particle identification. Resolution was calculated using a loose mask in Dynamo; final mask-corrected FSC was calculated in Relion using a two-pixel extended mask derived from the low-pass filtered 1SVD PDB structure (Extended Data Figure 6). For calculating resolution according to number of particles included, particles were first sorted by cross correlation to the final half-map, and the best-correlating particles were used according to the target percentage.

The general subtomogram averaging workflow was based upon a previous high-resolution approach from the Briggs group.^43^ Following particle identification, subtomograms were re-extracted from novaCTF weighted back projection tomograms. Carboxysomes were divided into half-sets, and subtomogram alignment and averaging was performed by progressively moving a lowpass filter to the resolution at which the FSC crossed 0.143 in the previous iteration.^41^ The binning and search space were decreased as resolution improved, and C4 symmetry was imposed after searches were restricted to less than 90-degree rotations. The best 80% of particles according to cross-correlation were used to generate the reference for the next iteration, but all particles were aligned for use in ultrastructure analysis. When the resolution ceased to improve by at least 1 Fourier pixel per iteration, the half-maps were sent to Relion for post-processing and final resolution calculations.^42^ Rigid body docking was performed in UCSF Chimera.^44^

Subtomogram averages of fibrils, monomer Rubisco, and outer layer Rubisco were calculated in Dynamo using the orientations and positions from the full-dataset alignment, C1 symmetry and no masking (see Data Analysis for classification approach). The outer layer analysis contained sufficient particles to average half-sets and perform post-processing in Relion, while the fibril average was computed from all particles and low-pass filtered to 1 nm due to particle number constraints. Difference maps were calculated by subtracting the average of unbound, non-outer layer Rubisco from the average of choice, and overlaid in Chimera using a 5 Å minimum size.

### Data analysis

Sixty-two tomograms containing 139 carboxysomes were collected, containing 32,930 identified Rubisco complexes. All of these data were used for subtomogram averaging. Further data analysis was only performed on CBs wholly within the field of view in all tilts, and with defoci allowing clear visual contrast (typically −2 μm or more), for a subset of 26,224 particles in 107 carboxysomes.

Carboxysome volume calculations utilize a previous observation that Rubisco forms an outer layer immediately adjacent to the shell.^14,17^ This behavior allows treatment of the Rubisco as a sphere in an accessible volume calculation. We defined the outer layer of Rubisco as the positions used in a convex hull calculation, and calculated the approximate volume contained by calculating the volume enclosed by each three-Rubisco facet plus a fourth reference point (a non-hull Rubisco position). Total Rubisco concentration was then calculated by converting Rubisco complexes per m^3^ to μM.

All data analysis calculations consider the D4 symmetry of the Rubisco complex. Dynamo uses a ZXZ’ Euler angle convention. Subtomogram averaging was performed to place the C4 axis of the complex along the Z axis in Dynamo, simplifying the handling of symmetry in the analysis. Only the Rubisco-Rubisco twist calculation considers the final azimuthal rotation (Z’); all other calculations only consider the orientation of the Rubisco C4 axis, using the orientation vector defined by the ZX Euler angles. Angles between Rubisco complexes are calculated using the formula

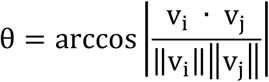

where v_i_ and v_j_ are the vectors defined by the C4 axis of two Rubisco complexes, and the absolute value of the dot product accounts for the identical head and tail due to the D4 symmetry. For calculations comparing the C4 Rubisco axis and the shell normal vector, the shell normal vector was calculated using a plane fit to the nearest facets in the convex hull calculation.

To search for Rubisco fibrils, we looped through every Rubisco centroid and set a search point 45 pixels along the C4 axis in both directions. Rubisco complexes are considered bound to the reference Rubisco if they have centers within 22.5 pixels of the search point and a C4 axis orientation differing less than 25 degrees from the reference. A Rubisco fibril is considered as inside a lattice if it is surrounded by 6 other fibrils within 70 bin2 pixels. A Rubisco is considered as being in the outer layer if it participates in the convex hull calculation.

Because the outer layer of Rubisco has location-specific properties, outer layer Rubisco complexes are not counted in fibril length even if they appear to participate. Outer layer Rubisco complexes are also excluded from the nearest-neighbor alignment analysis (Extended Data Figure 3). The nearest-neighbor analysis does not use total Rubisco concentration, but rather scales relative concentration of non-outer layer Rubisco (calculated as total inner Rubisco complexes divided by the CB volume).

The lattice visualization was performed in THREE.js. Rubisco alone or in a two-member fibril was made transparent, and the diameter was decreased to visually emphasize the pattern.

The collective alignment of Rubisco in the CB was estimated by calculating the second order tensor, Q_ij_, where v is the orientation vector of a Rubisco complex and δ is the Kronecker delta.^21^

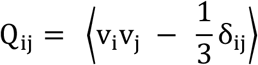

The scalar, S_ij_, is calculated from the largest eigenvalue of the tensor. It has a value range of 0 to 1, and the boundary between isomorphic and nematic phases occurs around 0.3-0.4.

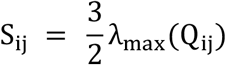

For means and errors presented in the text, all histograms were visualized. Distributions appearing to be Gaussian and with no scientific reason to assume otherwise are presented as a Gaussian mean and standard deviation. The bend angle of the Rubisco fibril, which shows a long-tailed distribution, is presented as the mode and full width half maximum. Plots showing histograms of angle orientations also display a line for random axis orientations, plotted with the same histogram binning and symmetry consideration.^45^

## EXTENDED DATA

**Extended Figure 1:**
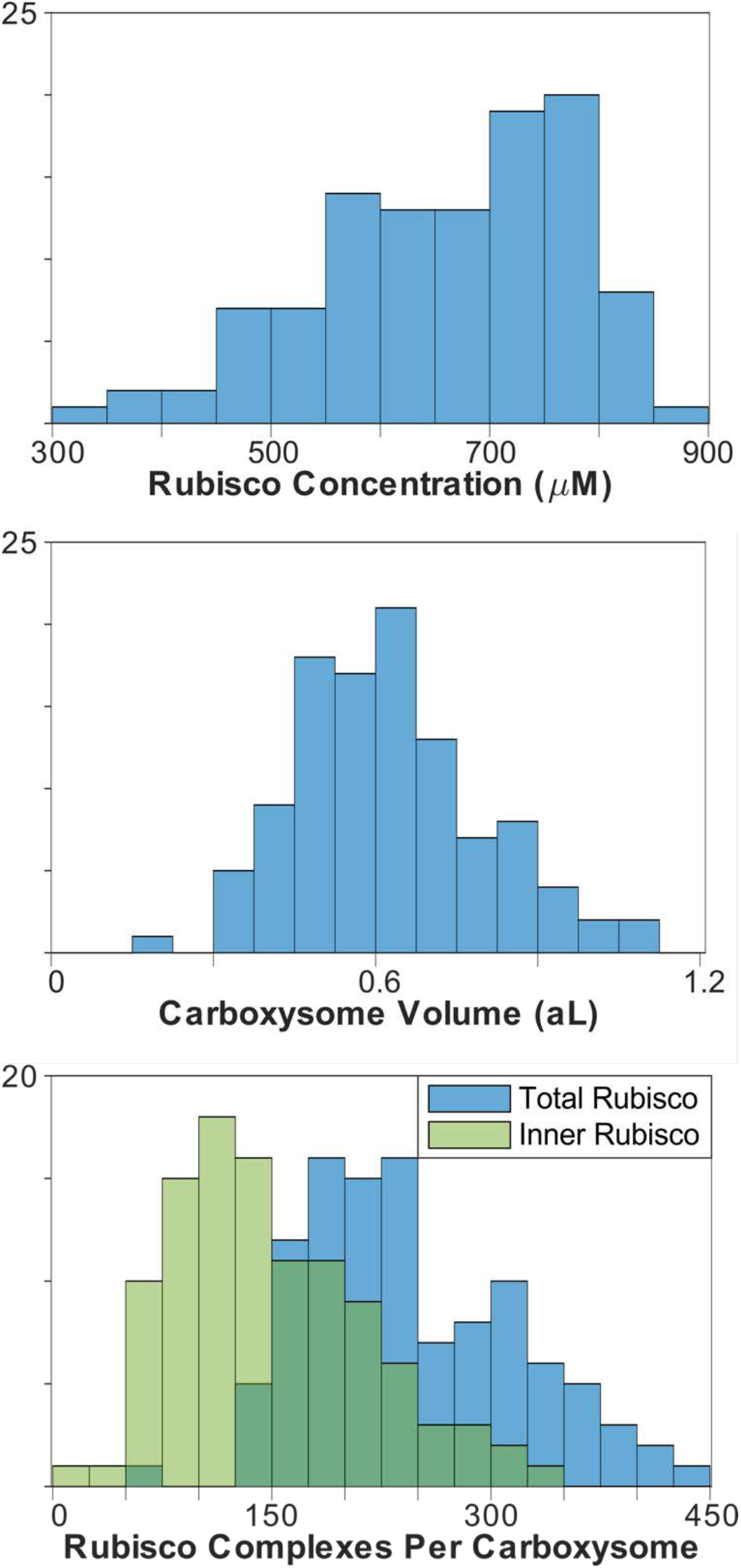
Carboxysome parameters. Top: Rubisco concentration per carboxysome. Middle: volume per carboxysome. Bottom: Rubisco complexes per carboxysome. Inner Rubisco excludes shell-adjacent Rubisco but does not adjust volume estimate.

**Extended Figure 2:**
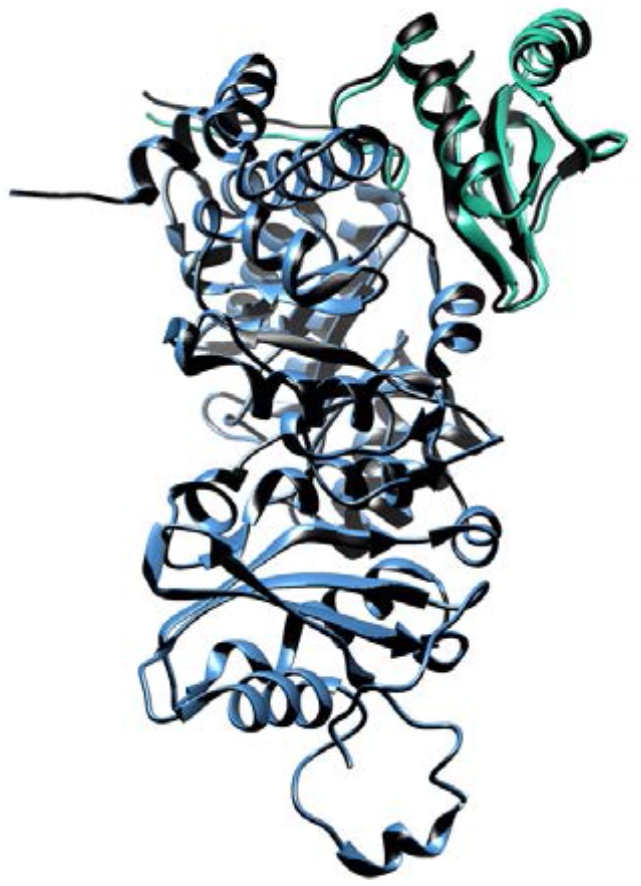
Orientation of the Rubisco subunits *in situ*. Black: 1SVD crystal structure. Color: large (blue) and small (green) subunits docked independently in our map.

**Extended Figure 3:**
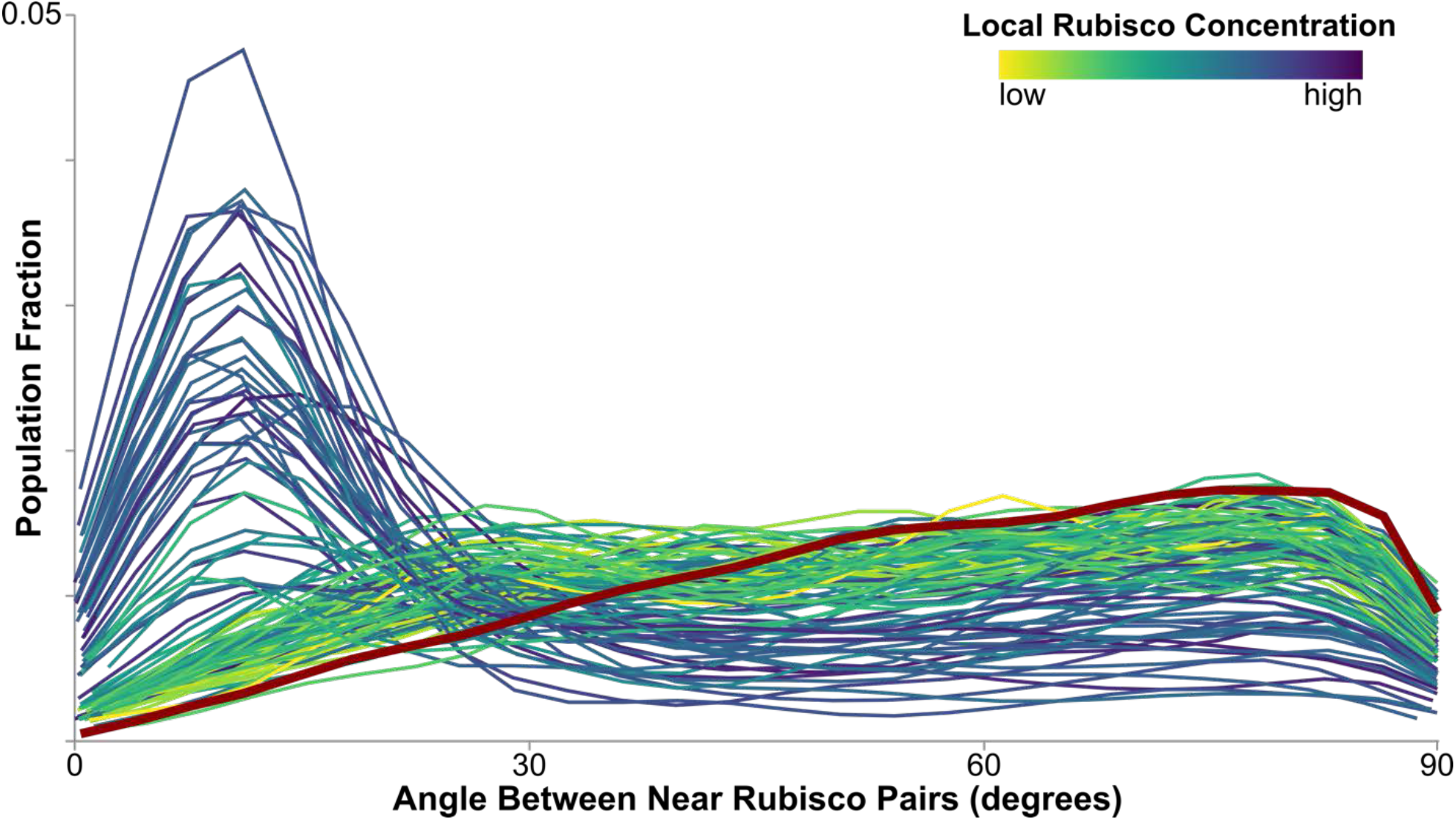
Histogram tracings of nearest-neighbor angle alignment inside carboxysomes. At low Rubisco concentrations, nearest neighbors have random orientation (red line). As concentration increases, Rubisco begins to align at a low nearest-neighbor angle.

**Extended Figure 4:**
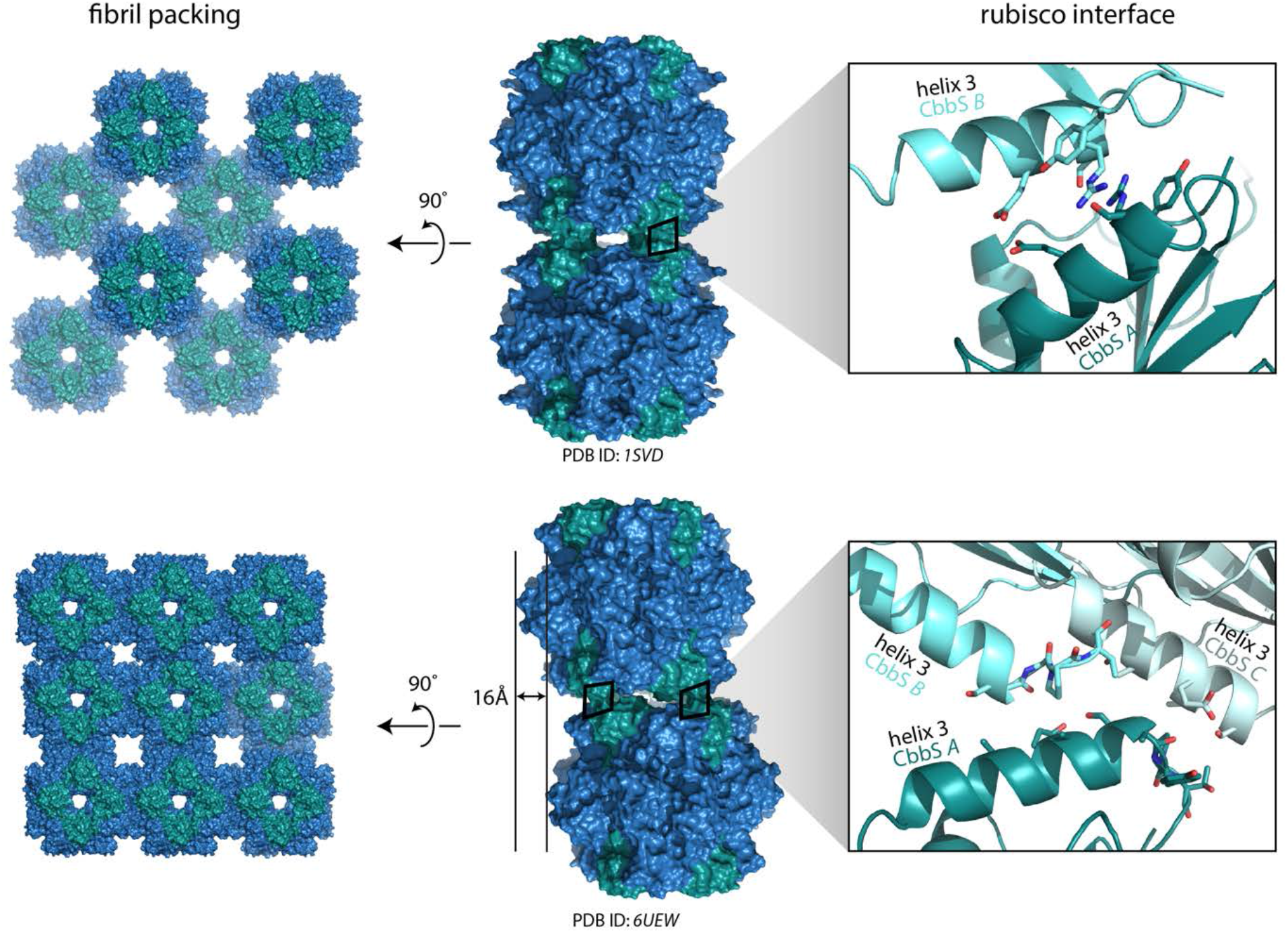
Longitudinal Rubisco fibril interfaces in crystal structures of Form 1A Rubisco (*H. neapolitanus*) from the Protein Data Bank. Blue, large subunit; green, small subunit. Both interactions are mediated by small subunit helix 3, but neither shows a specific, strong interaction between side chains.

**Extended Figure 5:**
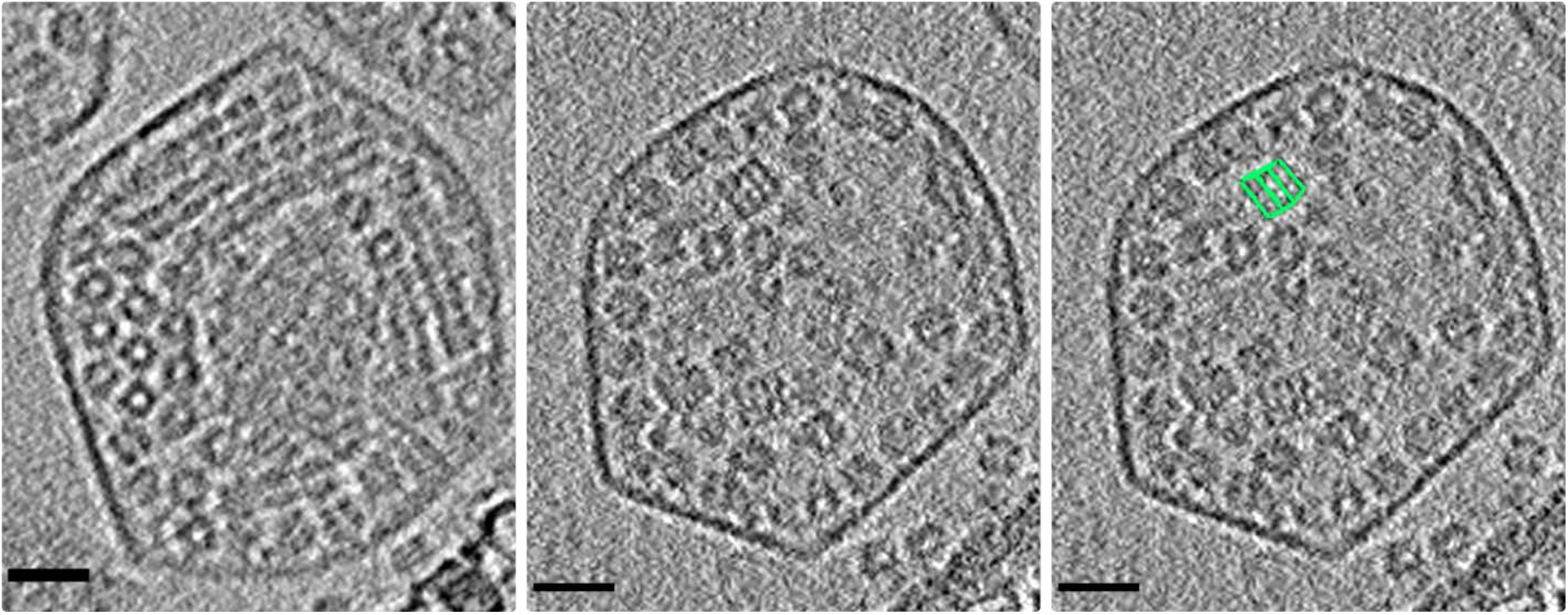
heterogeneity in carboxysome contents. Left: an 8.6 nm orthoslice through a carboxysome with fibrils arranged along the shell, rather than the standard central lattice. The center has a dense, poorly resolved area that may contain more disordered peptide. Center: a 4.3 nm orthoslice through a carboxysome containing two visible large non-Rubisco complexes. Right: one complex segmented in green. Scale bar 20 nm.

**Extended Figure 6:**
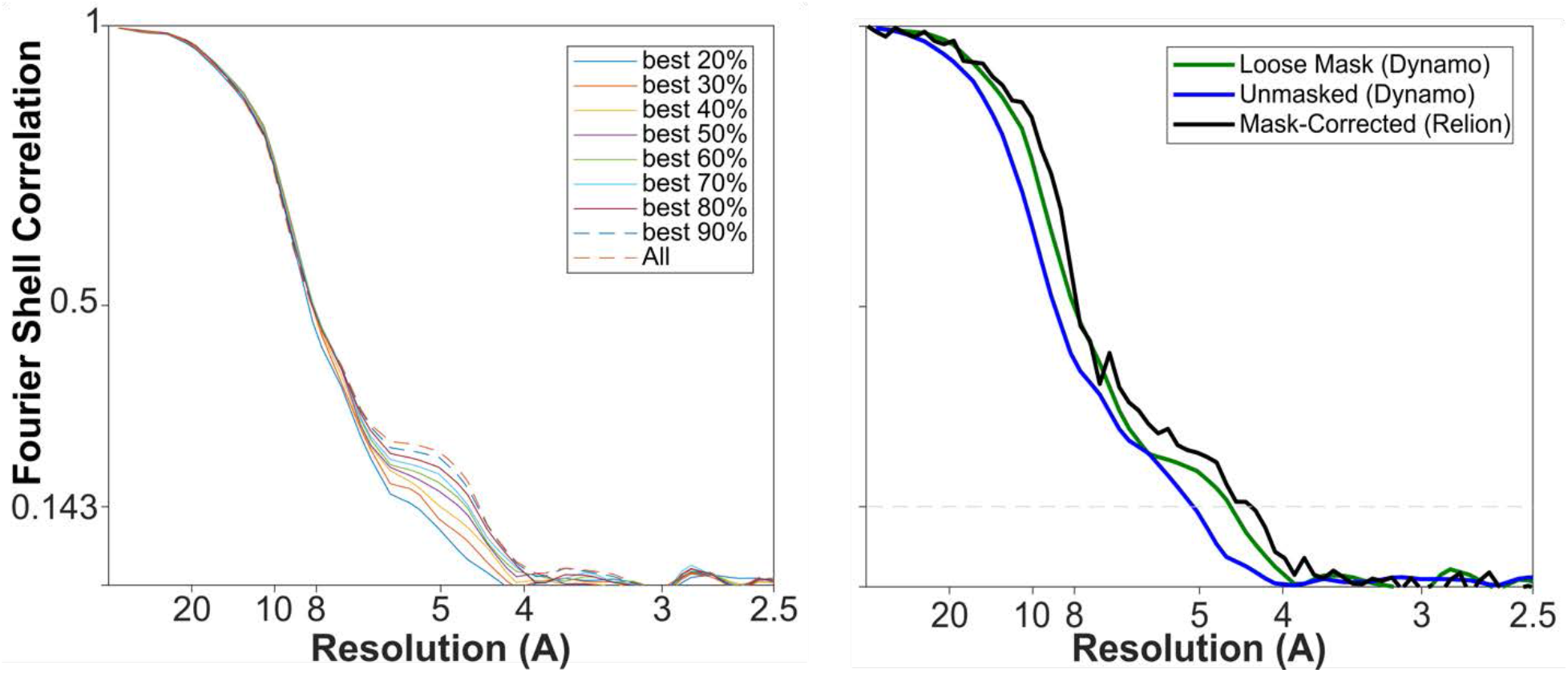
Left: Effect of particles on resolution. Removing the worst-correlating fraction of particles either decreases or does not affect resolution. Right: Fourier Shell Correlation for the final Rubisco subtomogram average. Unmasked FSC, blue; loose masked FSC, green; mask-corrected FSC, black. The resolution at 0.143 (grey dashed line) is 4.5 Å.

